# Social family structure and biogeography contribute to genomic divergence and cryptic speciation in the only eusocial beetle species, *Austroplatypus incompertus* (Curculionidae: Platypodinae)

**DOI:** 10.1101/2024.11.01.621599

**Authors:** James R. M. Bickerstaff, Bjarte H. Jordal, Markus Riegler

**Author notes:** Corresponding authors: James Bickerstaff; Markus Riegler.

## Abstract

Eusociality in insects has arisen multiple times independently in Hymenoptera (bees, wasps, ants), Blattodea (termites) and Coleoptera (beetles). In Hymenoptera and Blattodea, the evolution of eusociality led to massive species proliferation. In the hyperdiverse Coleoptera, eusociality evolved only once, in the ancient Australian ambrosia beetle species *Austroplatypus incompertus* (Curculionidae: Platypodinae). This species occurs in mesic eucalypt forests of eastern Australia, from Victoria to northern New South Wales. Based on few individuals collected from the southern and northern edges of the species’ distribution it was initially described as two distinct species; however, the names were later synonymised as no morphological differences were found in analyses of more specimens. Recent mitochondrial haplotype analyses revealed substantial latitudinal divergence across the distribution of *A. incompertus*. To address this apparent disparity between morphological and molecular data, we sequenced and analysed a SNP panel of over 6,656 biallelic markers from 187 individuals of 11 sites across 1000 km of this species’ range. Our data indicate that eusocial demographic processes such as low colony establishment success rate, limited dispersal and reliance on few reproductive individuals, together with substantial habitat fragmentation contributed to the population genetic structure of this species. We further identified that the Hunter Valley biogeographic barrier split the species into two distinct clades, with both clades in secondary close contact on the Barrington Tops plateau without any discernible admixture. Our results support the resurrection of a second species of *Austroplatypus* which has important consequences for the evolution eusociality in Coleoptera and the systematics of Platypodinae.

## Introduction

Eusociality, as the highest level of sociality in animals, is characterised by the division of reproductive and non-reproductive castes, cooperative brood care, and overlapping generations within a colony (Crespi & Yanega, 1995). In insects it has independently evolved in 15 orders multiple times and is associated with species radiations in the Hymenoptera (ants, bees and wasps) and Blattodea (termites) (Wilson & Hölldobler, 2005). Conversely, in other insect lineages the evolution of development of eusociality is rarer, such as in gall-forming aphids and thrips (Crespi, 1992; Stern, 1994). Social behaviours have also been described in the Coleoptera, with many ambrosia beetle species (Curculionidae: Platypodinae and Scolytinae) engaging in subsocial lifestyles such as colonies with overlapping generations and parental care of brood (Kasson et al., 2016; Kirkendall et al., 2015; Milbrath et al., 2024). Facultative eusociality has been demonstrated in four ambrosia beetle species; *Xyleborinus saxesenii* (Ratzeburg), *Xyelborus affinis* Eichhoff and *Trachyostus ghanaensis* (Schedl), whereby adults may refrain from reproduction (Biedermann, 2020; Biedermann et al., 2011; Kirkendall et al., 2015). However, true obligate eusociality, wherein there is a clear division of reproductive labour has only been demonstrated for one species, the Platypodinae ambrosia beetle *Austroplatypus incompertus* (Schedl) endemic to eastern Australia (Kent & Simpson 1992; Smith et al. 2018).

*Austroplatypus* is an ancient, roughly 50 Ma, monotypic genus that is sister to all other members of the Platypodini tribe (Jordal, 2015). It is younger than eusocial termites (Termitidae), wasps (Vespoidae) and ants (Formicoidea) (Jouault et al., 2021; Peters et al., 2017) but likely older than bees (Apidae) (Cardinal & Danforth, 2011; Peters et al., 2017; Schwarz et al., 2007). In contrast to the hymenopteran eusocial lineages, *Austroplatypus incompertus* has a diplodiploid sex determination system (Smith et al., 2009). It is primitively eusocial as no morphological differences exist between queens and workers (Kent, 2010; Smith et al., 2018). Colonies, also known as galleries, consist of a singly inseminated foundress queen and unmated daughter workers who do not leave the gallery, while other adult daughters and all adult sons leave (Smith 2018). The foundress copulates once and then initiates gallery formation without any further mating events, and sires have not been observed in colonies. No inbreeding events have been detected in families (Smith 2018) and as daughter workers remain unmated, the gallery is destined to collapse should the queen die. The foundress and her daughters construct galleries in living trees of stringybark *Eucalyptus* species distributed in discontinuous mesic forests along eastern Australia, from south-eastern Victoria to northern New South Wales (NSW) (Bickerstaff et al., 2020; Kent, 2008; Smith et al., 2018). Larvae and worker daughters contribute to gallery construction and the maintenance of symbiotic ambrosia fungus gardens (Beaver, 2000; Harris et al., 1976). The galleries are long lived and have been found to be maintained for up to 36 years (Harris et al., 1976; Smith et al., 2018). Young colonies retain daughter workers to assist in gallery maintenance, and as colonies age the proportion of dispersing daughters increases (Smith et al., 2018). Microsatellite analyses uncovered that relatedness between galleries rapidly decays across 500 m, suggesting that these beetles have a low dispersal capacity (Smith, 2013; Smith et al., 2018). Previous mitochondrial haplotype characterisation of three populations by Smith (2013) identified that the northernmost analysed population, Kerewong, NSW, had a pairwise divergence of 10% in the mitochondrial *cytochrome oxidase I* (COI) gene compared to other populations further south. Such strong social family structure detected at landscape and biogeographic scales are expected to affect overall population genetic structure and genomic divergence across the distribution of a species (Mikhailova et al., 2024).

Demographic factors and life history traitors strongly contribute to population genetic structure and species differentiation (Bickerstaff et al., 2023), and eusocial species are fantastic candidates to explore this (Brito et al., 2014; Landaverde-González et al., 2017). Common population genetic characteristics of eusocial species include low heterozygosity and strong population differentiation (Mikhailova et al., 2024), as reproduction eusocial colonies is typically performed by a single or few reproducing individuals. This has been observed in populations of the eusocial naked mole rat, *Heterocephalus glaber*, (Chau et al., 2018; Ingram et al., 2015; Zemlemerova et al., 2021) and eusocial subterranean termites (Bankhead-Dronnet et al., 2015; Dronnet et al., 2004, 2005; Fougeyrollas et al., 2018). In the case of haplodiploid eusocial Hymenoptera, low heterozygosity and effective population sizes, and strong population differentiation is also typical (Brito et al., 2014; Kahnt et al., 2014; Landaverde-González et al., 2017; Schenau & Jha, 2017; Seppä et al., 2009). Reduced gene flow between populations can increase the risk of inbreeding depression in these species (Grozinger & Zayed, 2020; López-Uribe et al., 2017; Lozier & Zayed, 2017), but is balanced in species with males that are capable of long-distance dispersal (Chapman et al., 2018; Wolf et al., 2012). However, most population genetic studies conducted on eusocial insects have focussed on the Hymenoptera and Blattodea, which may not be wholly representative given that eusociality occurs in other lineages, such as beetles and even vertebrate mole rats (Chau et al., 2018; Ingram et al., 2015; Zemlemerova et al., 2021).

Geographic processes can also contribute to population genetic structuring and further confound the interpretation of species boundaries (Hausdorf & Hennig, 2020). The spatial spread of organisms can limit the admixture of individuals across populations, leading to genotype fixation within populations, genetic drift and a loss of heterozygosity. With individuals mating only within population clusters and not fully admixing throughout the species range, individual genetic distances can become spatially autocorrelated, a scenario commonly referred to as isolation by distance (IBD) (Bradburd et al., 2018; Flanagan et al., 2021; Prunier et al., 2017). IBD can typically arise when the dispersal of taxa is hindered (but not totally obstructed) by geographic features and is mostly due to large pairwise distances. Impacts of IBD have been well documented and are known to inflate genetic distances between individuals and populations (Brito et al., 2014; Chapman et al., 2018; Toczydlowski & Waller, 2019) and can even lead to allopatric speciation events (Chan et al., 2020; Oberski et al., 2018; Rix & Harvey, 2012).

Eastern Australia is a well-studied natural laboratory for investigating the impacts of biogeography on population genomics. The Great Dividing Range (GDR) is a chain of mountain ranges and spurs that span eastern Australia in a north-south direction and provides habitat to pockets of mesic forest which are separated by large valleys and open woodlands and grasslands. The expanse of these mesic forest habitats has changed greatly throughout the geological history of Australia, with eastern Australia dominated by mesic forests throughout the Cretaceous and Paleogene epochs (145-23 Mya). These originally large forested areas began to contract and fragment roughly 65 Mya due to shifting rainfall patterns and aridification caused by moving plate tectonics and regional climatic warming (Bryant & Krosch, 2016). The aridification has reduced closed forest habitats throughout the Miocene-Pliocene epochs and resulted in the expansion of open dry-sclerophyll woodlands and grasslands. These dry areas are major biogeographic barriers to the dispersal for species adapted to the mesic habitat (Bryant & Krosch, 2016; Chapple et al., 2011; Flores-Rentería et al., 2021), and in many mesic species, phylogenetic divergences were formed by biogeographic barriers to dispersal (Bryant & Fuller, 2014; Chapple et al., 2011; Gunter et al., 2019; Pepper et al., 2014, 2018). Similar patterns are also seen in taxa with specialised host associations, such as specialised herbivores and pollinators. For example, the population structure of a fig psylloid, *Mycopsylla fici*, is characterised by high levels of population fragmentation and IBD along the GDR corresponding with biogeographic barriers (Fromont et al., 2017). Biogeographic barriers can further reinforce population structuring imposed by host distributions. Population structure in the fig pollinating wasp, *Pleistodontes nigriventris*, is structured by the host which only occurs in two widely separated rainforest blocks in eastern Australia (Cooper et al., 2020). Conversely, higher gene flow across barriers was observed between populations of *Pleistodontes imperialis* whose host tree has a wider, more connected distribution; while its parasitoid, *Sycoscapter sp.,* was identified to be panmictic throughout the entire distribution of its host with no impact of biogeographic barriers limiting gene flow (Sutton et al., 2016).

In this study we explored the interplay of social family structure and biogeography on the population genomics and structuring of the eusocial *A. incompertus*. The taxonomic history of the name *A. icompertus* has been tumultuous. This species was originally described as two, *Platypus incompertus* Schedl) (holotype female: NSW: Eden, 23.x.1953) and *P. incostatus* Schedl (holotype male: NSW: Dorrigo, 23.iii.1954), based on very few individuals from the southern and northern limits of this species’ distribution The distinction between these two species were made by incorrect inferences of sexually dimorphic traits such as mycetangia and elytral structure (Kent 2010). Browne (1971) erected the genus *Austroplatypus* and transferred *P. incompertus* to this new genus. Later, the names *P. incostatus* and *A. incompertus* were synonymised as the only observed difference between populations sampled throughout the range was body size variation along a latitudinal cline, possibly following Bergmann’s rule (Kent 2010). Given the importance of *A. incompertus* to the evolution of eusociality in Coleoptera, the fluctuating taxonomic history of this species and findings of mitochondrial divergence, we investigated the population structure and genetic diversity of *A. incompertus* using genome-wide Single Nucleotide Polymorphism (SNP) and mitochondrial markers. With these data we identified how eusociality, demographic and biogeographic processes contributed to on-going genomic divergence leading to speciation in a unique lineage of eusocial beetles.

## Methods

### Sample collection and DNA extraction

Individuals of *A. incompertus* were collected from 11 sites across 1000 km through most of the species range in eastern Australia (Figure 1) between 2008 and 2020. Brass gauze micro-cages (Smith et al., 2018) were fitted onto the exit of active galleries on stringybark *Eucalyptus* species in early March (before the start of emergence) and removed in late April (austral autumn). Adult live beetles were collected on regular site visits, immediately preserved in pure ethanol and then stored at –20 °C. Genomic DNA was extracted from 183 individuals across 125 galleries, with 1 to 3 individuals per gallery (Table S1), using a Qiagen DNEasy Blood and Tissue kit following a modified protocol. Beetles were ground with a pestle in a microcentrifuge tube containing 180 µL of Buffer ATL, after which 20 µL of Proteinase K was added, followed by a lysis incubation for 3 hours. The DNA was then eluted into 50 µL ultrapure water. All DNA extracts were assessed for quality and quantity with Nanodrop, Qubit fluorometry and gel electrophoresis.

**Figure 1.**
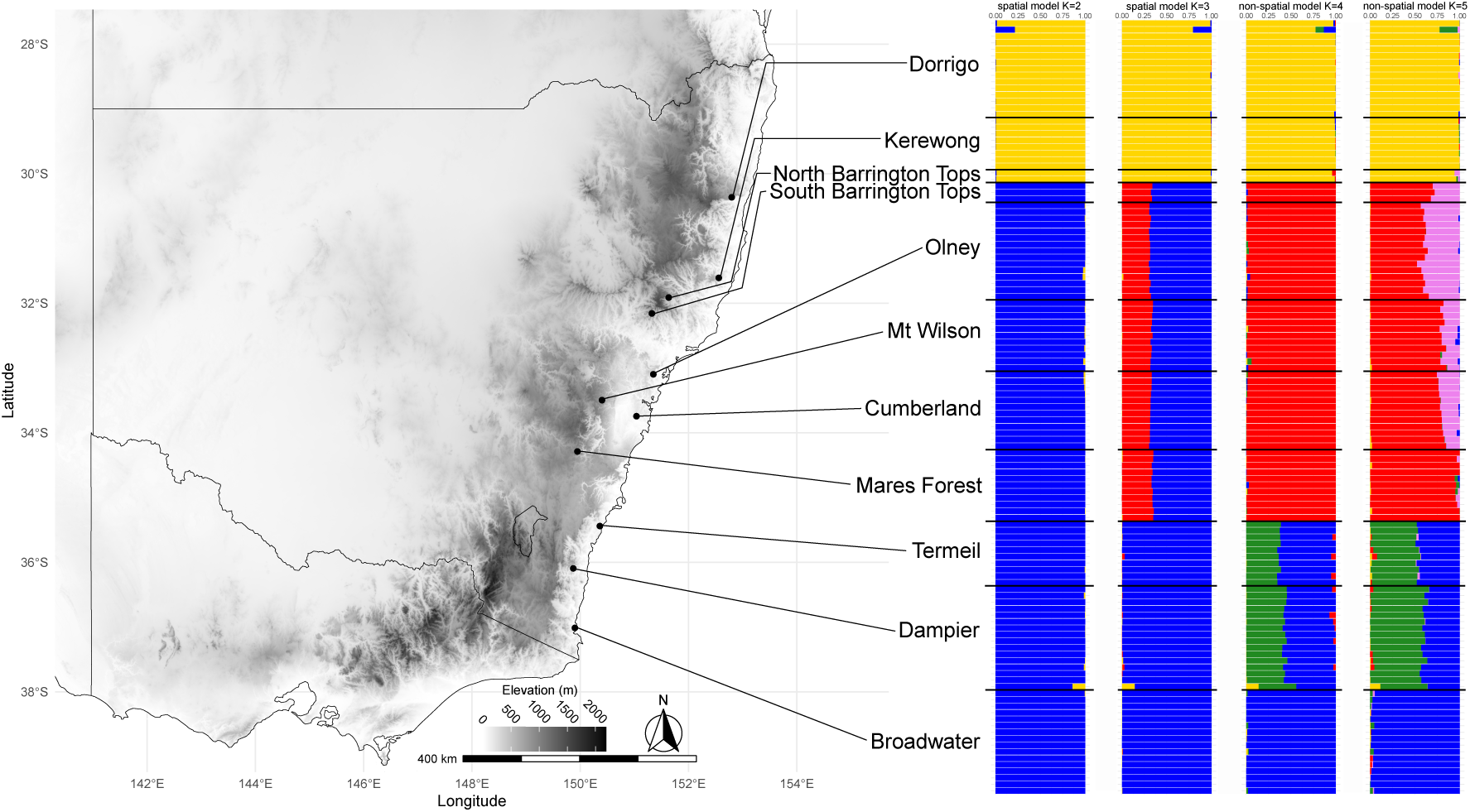
Map of southeastern Australian with the sites from where *Austroplatypus incompertus* individuals were collected. The admixture plots to the right show the conStruct population clustering with both spatial (K=2; K=3) and non-spatial models (K=4; K=5) across the two most optimal values of K. For all results each individual is partitioned in K-coloured segments representing shared genetic ancestry across clusters.

### Mitochondrial COI gene and reduced genome sequencing

A fragment of the mitochondrial COI gene was amplified for 35 specimens of *A. incompertus* which included at least three individuals from each site using the LCO1480/HCO2198 primer pair (Folmer et al., 1994). PCR reactions were performed in 10 µL containing 0.2 µL (20 nM) of each primer, 2 µL of 5x MyTaq Red Buffer, 0.2 µL of MyTaq DNA polymerase, 6.4 µL of PCR grade water and 1 µL of template DNA. Thermocycler conditions were six cycles of denaturation at 94 °C for 1.5 minutes, annealing at 45 °C for 1.5 minutes and extension at 72 °C for 1 minute, and then the annealing temperature was increased to 51 °C for further 35 cycles followed by a final extension at 72 °C for 5 minutes. Amplicons were Sanger sequenced in the forward direction. All chromatograms were assessed for quality, and chromatogram peaks and open reading frames for the presence of stop codons that could potentially indicate the presence of nuclear mitochondrial DNA (numt) (Bensasson et al., 2001).

DNA extracts of 183 samples were submitted to Diversity Arrays Technology (Canberra, Australia) for genotyping using DArTseq 1.0 complexity reduction technology. This approach utilises restriction enzymes, *EBPCR1* and *HpaII,* followed by fragment size selection to reduce genomic complexity prior to sequencing. Following enzymatic digestion and barcode ligation, libraries were then multiplexed and sequenced on an Illumina HiSeq 2500 platform. Single Nucleotide Polymorphisms (SNPs) were called using proprietary DArT analytical pipelines (DArTsoft14) by aligning sequence reads to the Dryophthorinae red palm weevil (*Rhynchophorus ferrugineus*, GCA_014462685.1) genome. This genome was chosen as Platypodinae (*Austroplatypus*) and Dryophthorinae (*Rhynchophorus*) are sister subfamilies within the Curculionidae, and as such SNPs called from this reference will reliably detect deeply diverging markers informative for the investigation of population structure and species boundaries.

### Mitochondrial COI phylogenetic and nuclear phylogenomic analyses

COI sequence alignment was performed using MAFFT v7.490 (Katoh, 2002) and reading frames were inspected by eye. We used maximum-likelihood (ML) and Bayesian approaches for phylogenetic estimation with the sequences partitioned by 1^st^, 2^nd^ and 3^rd^ codon positions, and trees were rooted with *Notoplatypus elongatus* Lea (HQ883688.1) and *Rhynchophorus ferrugineus* (Olivier) (MG051027). Maximum likelihood analyses were performed with IQ-TREE v2.2.0.5 (Minh et al., 2020), substitution models estimated using ModelFinder Plus and bootstrap support calculated with UFBoot (Hoang et al., 2018) using 1,000 replicates. Bayesian analyses were performed with BEAST2 v2.6.7 (Bouckaert et al., 2014) and the substitution model was selected using bModelTest (Bouckaert & Drummond, 2017). MCMC chains ran for 10,000,000 iterations with a burn-in of 10,000 and were logged every 10,000 chains. Convergence to stationarity and effective sample size values of all model parameters were assessed with TRACER v1.7.1 (Rambaut et al., 2018), and a maximum clade credibility tree with a 10% burn-in for all analysed datasets was inferred using TreeAnnotator v2.6.0. Following this, we calculated average nucleotide p-distances of the *COI* alignment between the northern, central and southern populations using MEGA 11 (Tamura et al., 2021) with 500 bootstraps, incorporating models of transition and transversion substitutions and uniform rates.

The SNP data were filtered using functions in the dartR v2.9.7 package (Mijangos et al., 2022). Monomorphic loci, SNPs with a reproducibility <99%, a read depth <7-fold and >50-fold, and loci with a call rate <90% were filtered. Additionally, a second dataset was made wherein only up to four individuals per population were retained. Using the gl2fasta function implemented in dartR, two alignments were made of both the full and reduced datasets. These alignments were then used for ML phylogenetics using IQ-TREE v2.3.2 (Minh et al., 2020) as above, except substitution models were estimated using the standard model selection, and ascertainment bias was corrected by invoking the +ASC command. Species trees were also constructed using the SNAPP package (Leaché et al., 2014) implemented in BEAST2. First, we sampled the mutation rate in the program, set the alpha to 1, beta to 250, lambda to 0.01 and coalescence rate to 10, and used default settings for all other parameters within beauti. Ten independent SNAPP analyses were performed, which were run for 10 million generations each, with 10% of the run discarded as burn-in. Convergence was assessed with TRACER v1.7.2 (Rambaut et al., 2018) and trees were summarised with treeannotator v2.6.4. All phylogenetic trees, for both *COI* and SNPs, were drawn with iTol (Letunic & Bork, 2024) and edited in Inkscape v2.

### Population clustering and structure

The population structure of *A. incompertus* was assessed with model-free K-means clustering approaches. First, SNPs that had a reproducibility of <95%, monomorphic loci and secondary SNPs (i.e. only the first SNP on a fragment was retained) were excluded. A minor allele frequency (MAF) threshold of >0.02 was applied, and all loci that deviated from Hardy Weinberg Equilibrium (HWE) were excluded. Paralogous loci (those with a Hamming distance between two sequences of >20%) were removed, and lastly individuals with a call rate <80% and loci with a call rate <75% were excluded. We first ran a Principal Components Analysis (PCA) to assess population structure with the dudi.pca function in the adegenet v2.1.10 package (Jombart, 2008). We then ran a cross-validation analysis to identify the Principal Components (PCs) that contributed most to the data structure, and the retained PCs were used in a Discriminant Analysis of Principal Components (DAPC) using the package adegenet (Jombart, 2008). We computed pairwise population level differentiation using the gl.fst.pop function in dartR with 1,000 bootstrap replicates for significance testing.

We explored the effect of spatial autocorrelation in our dataset as *A. incompertus* populations are clustered in habitat patches along eastern Australia, and our sampling throughout their range was discontinuous. Inferences of continuous genetic differentiation may be influenced by discreet geographical processes in that increasing geographic distances are linearly correlated with increasing genetic differentiation (Bradburd et al., 2018). As we are testing a potential allopatric speciation event between a set of lineages that follow a latitudinal cline, we needed to explore whether the observed genetic diversity between pairs are spatially autocorrelated (IBD). Therefore, we tested IBD by computing pairwise Provesti’s genetic distances, using the poppr v.2.9.6 package (Kamvar et al., 2014), and pairwise great circle distances in km, using the fields package, for all samples. A Mantel test was used to test the correlation between these pairwise datasets, using the function mantel in ecodist v2.1.3 (Goslee & Urban, 2007), with 100,000 permutations and 50,000 iterations for bootstrap support.

As IBD was found to impact our dataset, population genetic admixture was assessed with conStruct v.1.0.6, a model-based population clustering package (Bradburd et al., 2018). This method is similar to STRUCTURE-like analyses in that a Bayesian analysis based on K-means clustering methods is used to describe populations admixture in a given dataset. However, conStruct uses IBD to explain genetic variation with explicit spatial and genetic components to assign clusters. We modelled population structure of *A. incompertus* using both spatial and non-spatial conStruct analyses to explore whether population admixture is driven by IBD or an allopatric speciation event. First, we only retained one individual per gallery to avoid duplicate genotypes and then ran the models with a *K* of 1 to 5, with 10 replicates per value of *K,* for 100,000 iterations with a burn-in of 50,000 iterations. To evaluate the optimal value of *K*, we calculated layer contributions for each *K* value to assess which layers contribute most to total admixture. Spatial and non-spatial conStruct structure admixture plots were visualised for the top two optimal *K* values and illustrated with ggplot (Wickham et al., 2019) and edited in Inkscape v2.

### Population genetic diversity

To investigate whether the detected split in the population structure of *A. incompertus* at Barrington Tops (between North and South Barrington Top) is due to an allopatric vicariance event or secondary contact of the northern and southern clades, we explored population genetic diversity. As species migrate and expand their range, the genetic diversity and heterozygosity at the edge of the range is reduced due to genetic drift (Jaya et al., 2022). As such, should the genetic diversity and heterozygosity be lower in the Barrington Tops populations, this would indicate range expansions of the northern and southern clades leading to secondary contact. If, however, genetic diversity and heterozygosity are similar to other populations or higher in the Barrington Tops populations, this could suggest a vicariance event which led to the allopatric speciation of *A. incompertus* had occurred in this area. To explicitly test this, we assessed population genetic diversity in all individuals. First, SNPs with a <99%, monomorphic loci, loci with a call rate >95% and individuals with a call rate >90% were filtered out, for all three datasets. Heterozygosity and Shannon’s Diversity were estimated for all populations in all datasets using the gl.report.heterozygosity and gl.report.diversity functions in the dartR package (Mijangos et al., 2022). We then explored mutation-drift equilibrium by calculating Tajima’s *D* for all populations in all datasets, using scripts produced by Jaya et al. (2022). For this, excessive rare alleles that are consistent with range expansions are indicated by significant negative values, while population contractions are indicated by significant positive values. Lastly, an analysis of molecular variance (AMOVA) based on Euclidean genetic distances between samples within northern and central/southern clades were performed to assess hierarchical genetic structure between populations, galleries sampled and individuals within a population. Significance for all tests was assessed using 999 permutations using the poppr.amova function in the poppr package (Kamvar et al., 2014).

## Results

In total 6,656 biallelic SNP markers were called prior to filtering and dataset assignment for 187 samples, with only one sample not meeting the minimum sequence call threshold. The average read depth, or coverage, of loci was 36-fold, and the average reproducibility of loci was 99%. The average call rate of loci and across samples was 60%. The total number of loci used for analyses varied from 452 (phylogenetic analyses) to 1,080 (population structure analyses) following varying filtering parameters.

### Phylogenetic analyses of mitochondrial COI and genomic SNP data

For the mitochondrial *COI* gene, two strongly supported clades were identified in both Bayesian (Figure 2a) and ML (Figure S1) phylogenetic analyses. One clade consisted only of individuals of the northern populations, while the second clade consisted of individuals from the central and southern populations. Furthermore, central and southern populations clustered in separate subclades except for some southern Termeil samples. The average pairwise distances between northern and other populations (p-distance North vs Central = 0.1086, p-distance North vs South = 0.1084) were greater than the distance between central and southern populations (p-distance = 0.0299). Similar to the *COI* phylogenetic trees, two strongly supported clades were identified in the SNAPP Bayesian (Figure 2b) and ML (Figures S2 & S3) phylogenetic analyses of the genomic SNP data, with one clade consisting of individuals of northern populations, while the second clade consisted of individuals from central and southern populations; each resolved monophyletically and strongly supported.

**Figure 2.**
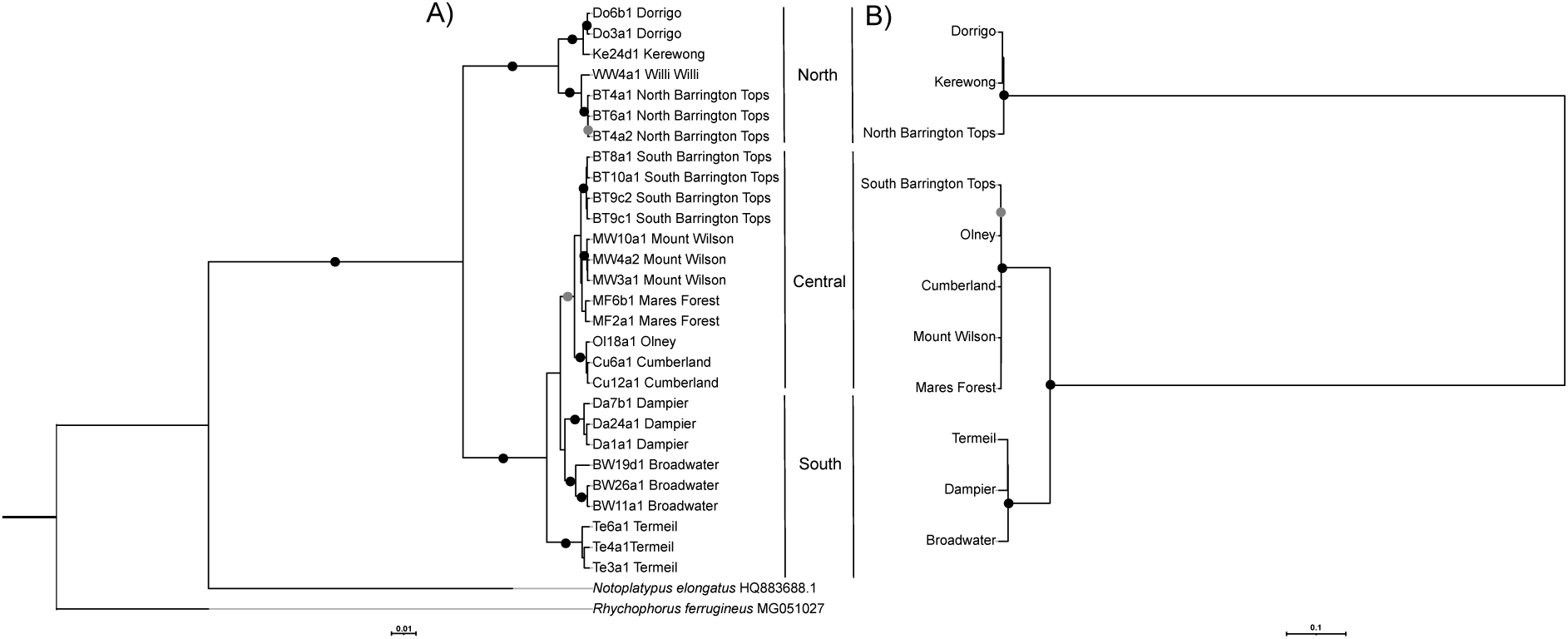
A) Bayesian phylogeny of *Austroplatypus incompertus* inferred from 654 bp of the mitochondrial *cytochrome oxidase I* (COI) gene. Support is provided at the nodes, with black dots indicating >0.95 posterior probability and grey dots indicating >0.85 posterior probability. B) Bayesian unrooted phylogeny of *A. incompertus* from the SNAPP Bayesian analysis inferred from 452 SNP markers. Support is given at the nodes, with black dots indicating >0.95 posterior probability and grey dots indicating >0.85 posterior probability.

### Spatial dynamics of Austroplatypus

Three clusters were identified with PCA (Figure 3a) and DAPC (Figure 3b), which included northern (Dorrigo, Kerewong and North Barrington Tops), central (South Barrington Tops, Olney, Mt Wilson, Cumberland and Mares Forest) and southern (Termeil, Dampier and Broadwater) populations, similar to what was identified in the phylogenetic analyses. Individuals across populations clustered tightly, to the exception of South Barrington Tops and Broadwater. Populations of *A. incompertus* were strongly differentiated in both northern and southern datasets (Figure 4). Despite the close geographic distance between Kerewong and North Barrington Tops (roughly 90 km) they were strongly differentiated from one another (F_st_ = 0.425). Mares Forest and South Barrington Tops were most distinct from the other populations within the central clade. Broadwater was strongly differentiated from Dampier and Termeil populations (F_st_ = 0.641 & 0.62, respectively).

**Figure 3.**
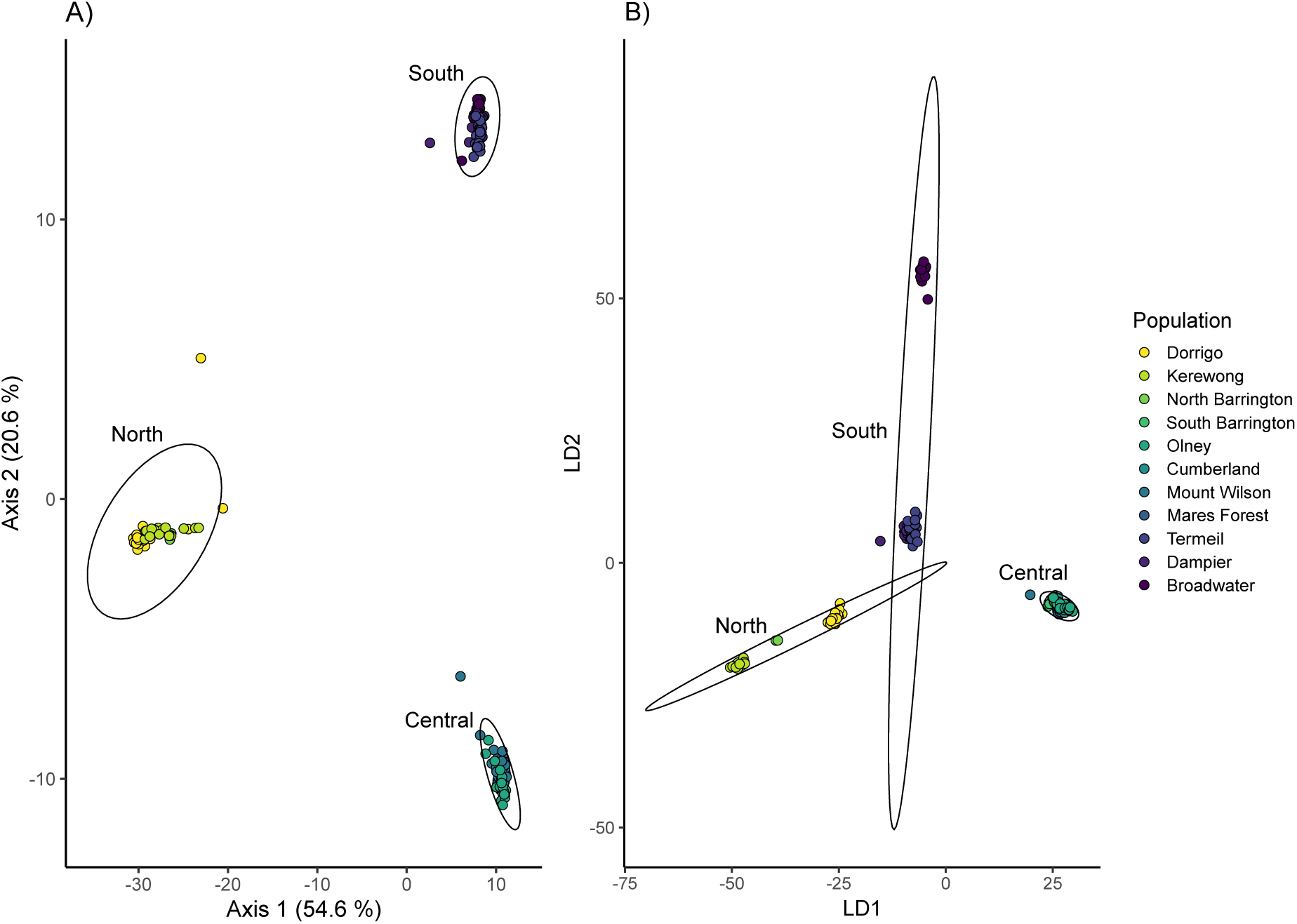
Both A) Principal Components Analysis (PCA) and B) Discriminant Analysis of Principal Components (DAPC) of nuclear SNP data identified three genetic groups corresponding with geographic clusters (denoted by ellipses) of *Austroplatypus incompertus*.

**Figure 4.**
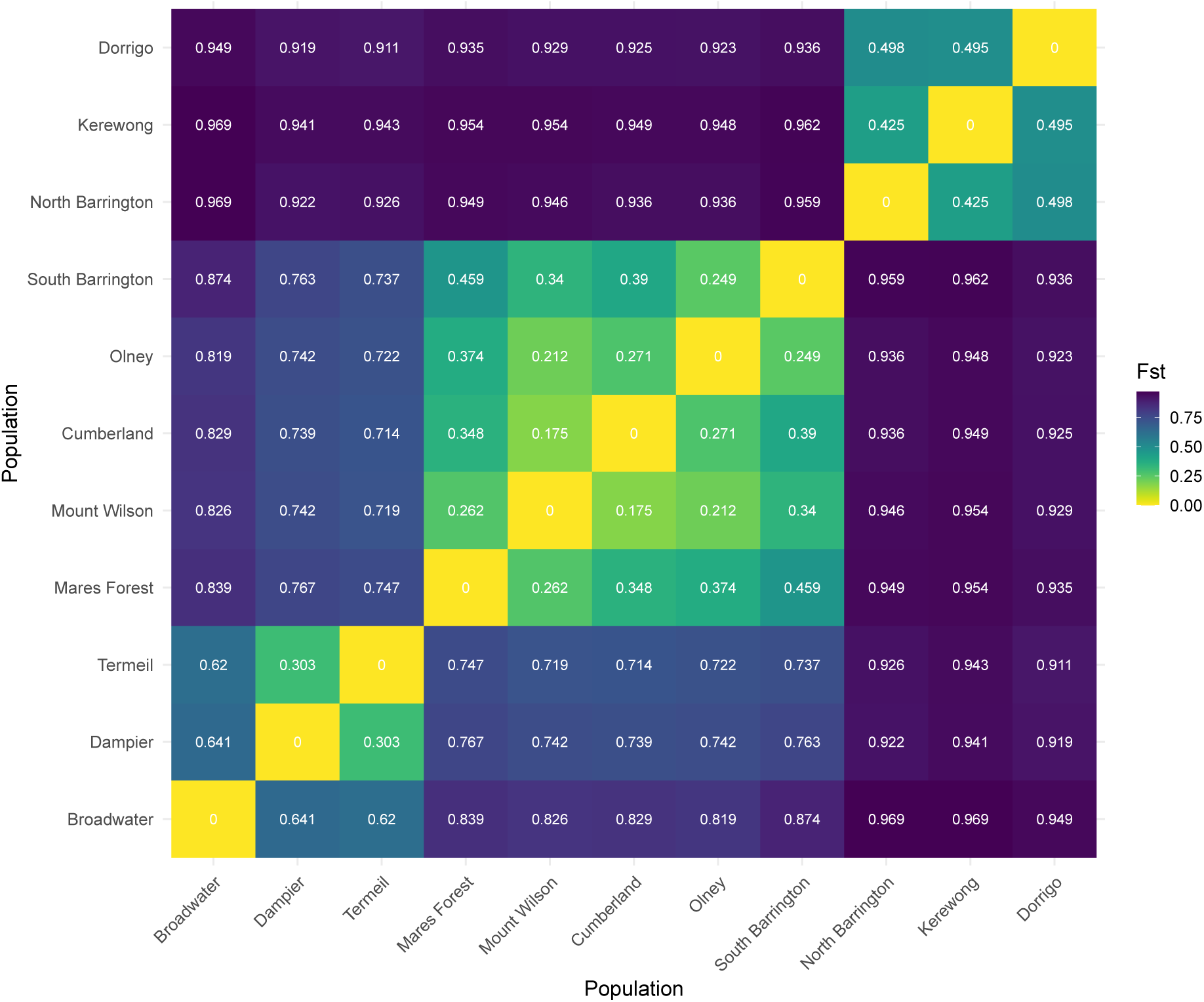
Pairwise heatmap showing Fst genetic differentiation among populations of *Austroplatypus incompertus*.

We also found strong support for spatially autocorrelated genetic distances and IBD (Mantel’s r = 0.7745, *p* < 0.001; Figure S4). Genetic clustering identified was similar to that of phylogenetic, PCA and DAPC analyses. The non-spatial conStruct model identified strong support for K = 4 and 5 while for the spatial model K = 2 and 3 were supported, by cross-validation. In the non-spatial model, as K increased from 4 to 5, additional genetic diversity and admixture were observed within the central populations. Conversely in the spatial model, two primary genetic clusters were identified, which included the northern population cluster and the combined central and southern population cluster. Increased genetic diversity and admixture were observed in central populations when K increased from 2 to 3. Admixture between clusters was only identified within and across central and southern populations in all models analysed (Figure 1). Additionally, the tested models did not identify gene flow between South and North Barrington Tops despite their close geographic proximity (<40km).

### Population genetics and variation

Across all datasets, we found that populations in Barrington Tops (South Barrington Tops and North Barrington Tops) had low heterozygosity and Shannon diversity compared to other populations. Olney, Cumberland, Dampier and Kerewong were highest in heterozygosity measures, and Olney, Cumberland, Dampier and Termeil had the highest Shannon diversity (Table 1). Measures of Tajima’s D was only supported for Dorrigo, which was negative suggesting an excess of rare alleles indicative of a potential recent population expansion (Table 1). The largest source of genetic differentiation was between populations (northern clade: 58%, σ = 67.98, *p* < 0.01; southern clade: 72%, σ = 202.47, *p* < 0.01). Relatedness between colonies within populations contributed little to total genetic differentiation (northern clade: 10%, σ = 12.21, *p* < 0.01; southern clade: 6%, σ = 17.12, *p* < 0.01).

**Table 1.**
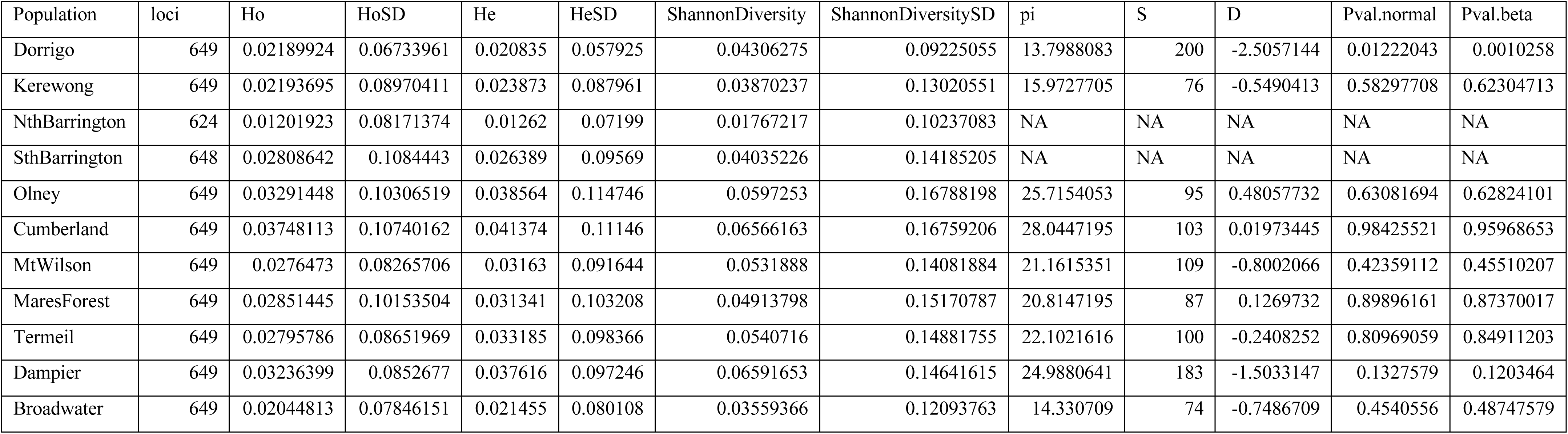
Genetic diversity indices of sampled *Austroplatypus incompertus* populations. Missing data indicate the average percentage of loci not sampled within that population. Ho: observed heterozygosity; He: expected heterozygosity; SD: standard deviation; pi: nucleotide diversity; S: number of segregating sites; D: Tajima’s D; Pval.normal: *p*-value calculated for Tajima’s D assuming a normal distribution; and Pval.beta: *p*-value calculated for Tajima’s D assuming a beta distribution.

## Discussion

Our study demonstrates that the evolution of eusociality in *Austroplatypus* is associated with ongoing population genetic divergence and contributes to cryptic speciation. Similar to other eusocial Hymenopteran (ants, bees and wasps) and Blattodea (termite) lineages, low heterozygosity, low gene flow between clusters and high Fst characterises the population genetic structure of this ambrosia beetle species. Genome-wide and mitochondrial *COI* markers consistently uncovered a deep divergence across populations of *A. incompertus*. Many of the 6,656 SNP markers were not shared by all populations (only 134 markers were shared), indicating a high proportion of private alleles within populations that could have arisen due to limited gene flow. Furthermore, biogeographic processes have further contributed to the ongoing divergence in this this species. In all clustering and phylogenetic analyses, including those that incorporate IBD tests, we consistently found that individuals from populations north of Barrington Tops formed a separate lineage distinct from populations south of Barrington Tops. Most importantly, though, we found individuals of the northern and southern lineages in North Barrington Tops and South Barrington Tops, respectively, with no detected gene flow between the two despite a distance of <40km between sampling sites. These results suggest that *A. incompertus* may be composed of two species. This finding has implications for the tribal level systematics of Platypodinae (Jordal, 2015; Kent, 2010) and the behavioural diversity and evolution of eusociality in Coleoptera.

### Eusociality drives population genetic structuring in Austroplatypus

*Austroplatypus* is characterised by low heterozygosity and high F_st_ throughout its range, similar to many other eusocial insect species (Brito et al., 2014; Kahnt et al., 2014; Landaverde-González et al., 2017; Schenau & Jha, 2017; Seppä et al., 2009). Results from AMOVA highlight that the variation in genetic diversity between galleries within a population contributes very little to the overall genetic structure of the species. We predict that eusocial colony structure, low dispersal capabilities and the spatial organisation within landscapes have contributed to low genetic diversity and high genetic divergence between populations in this species. *Austroplatypus* families consist of a singly and life-time inseminated foundress, permanent unmated daughter workers and immature full-sib offspring (Kent & Simpson 1982). In the early stages of colony activity, daughters are retained in the gallery while sibling males disperse whereas in later stages an increasing number of daughters disperse also (Smith et al. 2018). The ecological pressure of establishing a gallery in a live *Eucalyptus* tree is a costly, and potentially deadly (due to tree resin flow or predation), endeavour. Up to 90% of new galleries fail (Smith 2018) and successful, long-lived galleries require maintenance not only provided by the foundress but also her female offspring. Given this, genetic diversity may be bottlenecked due to the restricted number of reproductive foundress queens that are only inseminated once. Furthermore, the low dispersal capability of this species (Smith 2013) and the patchy distribution of galleries throughout their range despite, the presence of suitable hosts (Kent 1997, JB personal observations), has undoubtedly contributed to patterns of its genetic diversity and ongoing genomic divergence between populations.

### Australian biogeography contributes to speciation within Austroplatypus

The deep genetic divergence between *Austroplatypus* populations adds to a growing body of research that highlights the biogeographic effects on population genetic structuring and divergence of species in eastern Australia. Our analyses consistently identified breaks in gene flow between *Austroplatypus* populations, and to a lesser degree within geographic clusters. Three primary clades were identified: a northern clade (Dorrigo, Kerewong, North Barrington Tops), a central clade (South Barrington Tops, Olney, Mt Wilson, Cumberland, Mares Forest) and a southern clade (Termeil, Dampier, Broadwater). Population genetic structure of these clades aligned with two known biogeographic breaks: the Hunter Valley and the Southern Transition Zone south of the Blue Mountains (Bryant & Krosch, 2016). Ongoing gene flow was identified between central and southern clades while no gene flow was identified between northern and other clades. As central clade populations were identified to be more diverse than others, it is possible that *Austroplatypus* radiated northward and southward from the central region. Breaks in gene flow and phylogenetic patterns between populations of *Austroplatypus* are highly similar to other mesic forest adapted species which are constrained in their distribution by biogeographic barriers (Bryant & Fuller, 2014; Chapple et al., 2011; Gunter et al., 2019; Pepper et al., 2014, 2018).. Most of the genetic differentiation across populations was correlated with geographic distances between populations. The main isolating factor seems related to habitat fragmentation. *Austroplatypus* constructs galleries in stringybark eucalypts in mesic forests (Bickerstaff et al., 2020; Kent, 2008), and a reduction in suitable mesic forest has limited gene flow between isolated populations, resulting in the strong structuring identified across barriers identified herein.

Biogeographic barriers may have led to allopatric divergence and cryptic speciation within *Austroplatypus*. Individuals belonging to the northern clade and the central/southern clade were both collected on the Barrington Tops plateau, a locality to the north of the Hunter Valley biogeographic barrier. We identified no gene flow between these populations despite their close geographic proximity to one another. Interestingly, heterozygosity and Shannon diversity was lower in individuals sampled from the northern and southern areas of Barrington Tops compared to those of other populations, and this is indicative of a recent range expansion. Therefore, these lineages may be in secondary contact following their allopatric divergence due to aridification and formation of the Hunter Vallery biographic barrier. Such contact zones have been important for the identification of parapatric and allopatric speciation events (Brito et al., 2014; Moritz et al., 2018). Given that we detected no gene flow between the northern and central clades, this may suggest that *Austroplatypus* contains two distinct species. Therefore, the well-studied Hunter Valley biogeographic barrier (Bryant & Krosch, 2016) might have separated these two *Austroplatypus* clades through the aridification and reduction of closed mesic forest ecosystems some 10 Mya (Chapple et al., 2011). Further genotyping and morphometrics of individuals collected across the Barrington Tops region is needed to validate whether hybrids exist and whether the Hunter Valley was the allopatric driver of speciation.

### The “re-discovery” of a second eusocial beetle species

The discovery of a second *Austroplatypus* species is not just important for the taxonomic status of this species but also for tribal-level systematics within Platypodinae. In this study, we have sampled and sequenced individuals from sites in Dorrigo and close to Eden. The absence of gene flow between northern and central clade individuals in secondary contact in Barrington Tops support existence of two isolated lineages. As the northern clade of *Austroplatypus* may represent a distinct species, the taxonomic classification prior to Kent (2010) may need to be re-established. The holotype of *Platypus incostatus* (Schedl 1954) was represented by a male individual collected in Dorrigo, while the holotype of *A. incompertus* (Schedl 1953) was represented by a female individual collected in Eden in southern NSW. We, therefore, propose the establishment of *A. incostatus,* previously named *P. incostatus* by Schedl (1954), represented in our study as the northern clade. A second species in the genus may also support the erection of the Austroplatypodina sub-tribe, discussed further in Jordal (2015).

Our phylogenetic and phylogenomic evidence for the existence of two *Austroplatypus* species has important consequences for the understanding of the evolution of eusociality in beetles.

While research on the behaviour of *Austroplatypus* has focussed on the southern lineage (*A. incompertus*), i.e. in the Cumberland State Forest in Sydney (Kent & Simpson 1992, Smith 2018), and in Olney and Ourimbah State Forests on the Central Coast of NSW (Smith 2018), it is highly likely that colonies of the northern lineage (*A. incostatus*) are also eusocial given that they do not differ in any morphological, life-history and ecological features (Kent, 2008). Future studies should explore the behaviour, life history and ecology of *A. incostatus* to explicitly test whether they too are eusocial and whether any behavioural differences exist.

This will identify whether both species are eusocial and how species divergence has impacted eusocial behaviour and colony life history following speciation, which will be important for comparisons with other eusocial insect lineages. Furthermore, identification as to whether *A. incostatus* will allow for an accurate identification as to when eusociality evolved in Coleoptera. The lineage leading to *Austroplatypus,* and consequently the maximum age of the lineage of eusociality, is currently estimated to be 45-64 Myo (Jordal, 2015). Should only *A. incompertus* be eusocial, this would imply that the age of this behaviour is much younger, possibly 5 Ma (based on pairwise COI divergence).

The identification of a second eusocial beetle species also highlights a potential need for conservation action. Between August 2019 and February 2020, bushfires burned continuously across 10 million hectares (approx. 25 million acres), of which 80% was native forest (Saunders et al., 2021). Much of these bushfires burned through the distribution of both *Austroplatypus* species (Marsh et al., 2022). Given their low dispersal capability, low genetic diversity, and gene flow between populations, these recent catastrophic bushfires may have led to population decline in both species. Collections of these species were attempted by JRMB in 2022 in Broadwater and Termeil, sites from which individuals were collected for study in 2009 and 2016 and were devastated by the 2019-2020 bushfires. No active *Austroplatypus* galleries were found. Future studies should develop assessments of these species as to whether conservation listing is required to conserve and protect this unique eusocial beetle lineage.

### Conclusions

Herein, we identify that social structure and Australian biogeography contributes to ongoing genomic diversity of the world’s only eusocial beetle lineage, *Austroplatypus*. Our study has demonstrates that this mesic adapted and eucalypt specialist is not only dispersal limited but populations can persist despite low levels of heterozygosity and gene flow, similar to many other eusocial insect species (Bankhead-Dronnet et al., 2015; Dronnet et al., 2004, 2005; Fougeyrollas et al., 2018). By using multiple analytical approaches on these genetic data, we have identified a likely second species of *Austroplatypus*. Despite decades of changes in the taxonomic status of *A. incompertus* (Schedl 1953, 1954, Kent 2010), the integration of novel data and analytical techniques have yielded considerable new insights into the species boundaries and delimitation of this species and also is important for the systematics of Platypodinae. The identification of a second eusocial beetle species opens up new avenues in research to identify potential differences or similarities in how eusocial behaviours are expressed in these diverging populations. Our study presents an exciting new avenue in not only testing species boundaries, but also how population genomic research can impact our inference on species evolution and behaviour.

## Acknowledgements

We thank Shannon Smith and Robert Mueller for access to some of the specimens used in this study. We also thank Shannon Smith, Robert Mueller, Deborah Kent, Alf Britton and Scott Nacko for assistance with field collections, as well as Deborah Kent for the extensive advice on the life history, ecology and biology of *Austroplatypus*.

## Author contributions

JRMB, BHJ and MR conceptualised the research and designed the experiments. JRMB and MR collected the data, JRMB, BHJ and MR analysed and interpreted the data. JRMB wrote the manuscript with input of BHJ and MR. MR and JRMB were responsible for funding. All authors agreed to the submitted manuscript.

