## Supplementary Information for "Social family structure and biogeography contribute to genomic divergence and cryptic speciation in the only eusocial beetle species, *Austroplatypus incompertus* (Curculionidae: Platypodinae)"

**Supplementary information for:** Two for one – population genomics of the only eusocial beetle species, *Austroplatypus incompertus*, suggests the presence of a second species

**Authors:** James R. M. Bickerstaff[1, 2] *, Bjarte H. Jordal[3], Markus Riegler[1]*

[1] Hawkesbury Institute for the Environment, Western Sydney University, Locked Bag 1797, Penrith, NSW 2751, Australia

[2] Australian National Insect Collection, CSIRO, GPO Box 1700, Canberra, ACT 2601, Australia

[3] Museum of Natural History, University Museum of Bergen, University of Bergen, NO-5020 Bergen, Norway

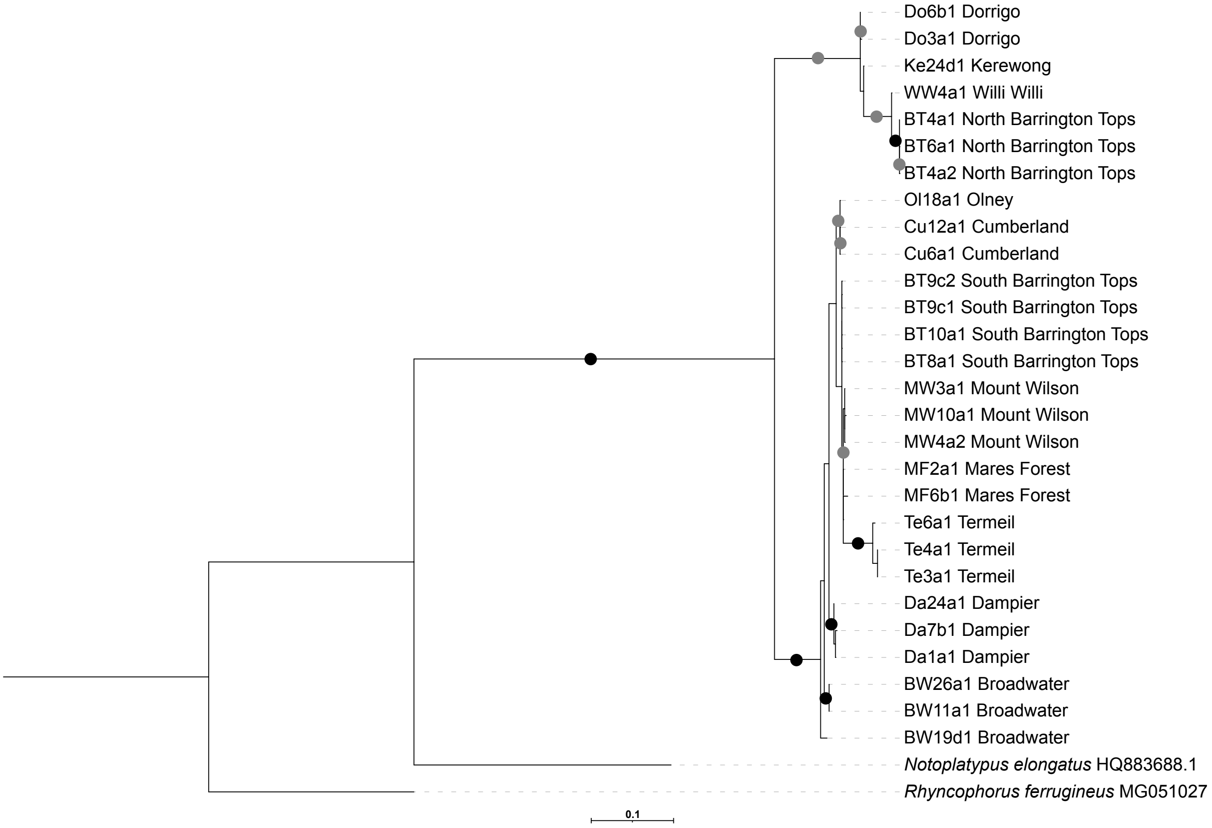


**Figure S1.** Maximum likelihood phylogeny inferred from 654bp of the *cytochrome oxidase I* gene. Support is given on the branches, with black dots indicated >95% bootstrap support and grey dots indicating between 85 – 94% bootstrap support. Branches with no dots indicate <84% support.

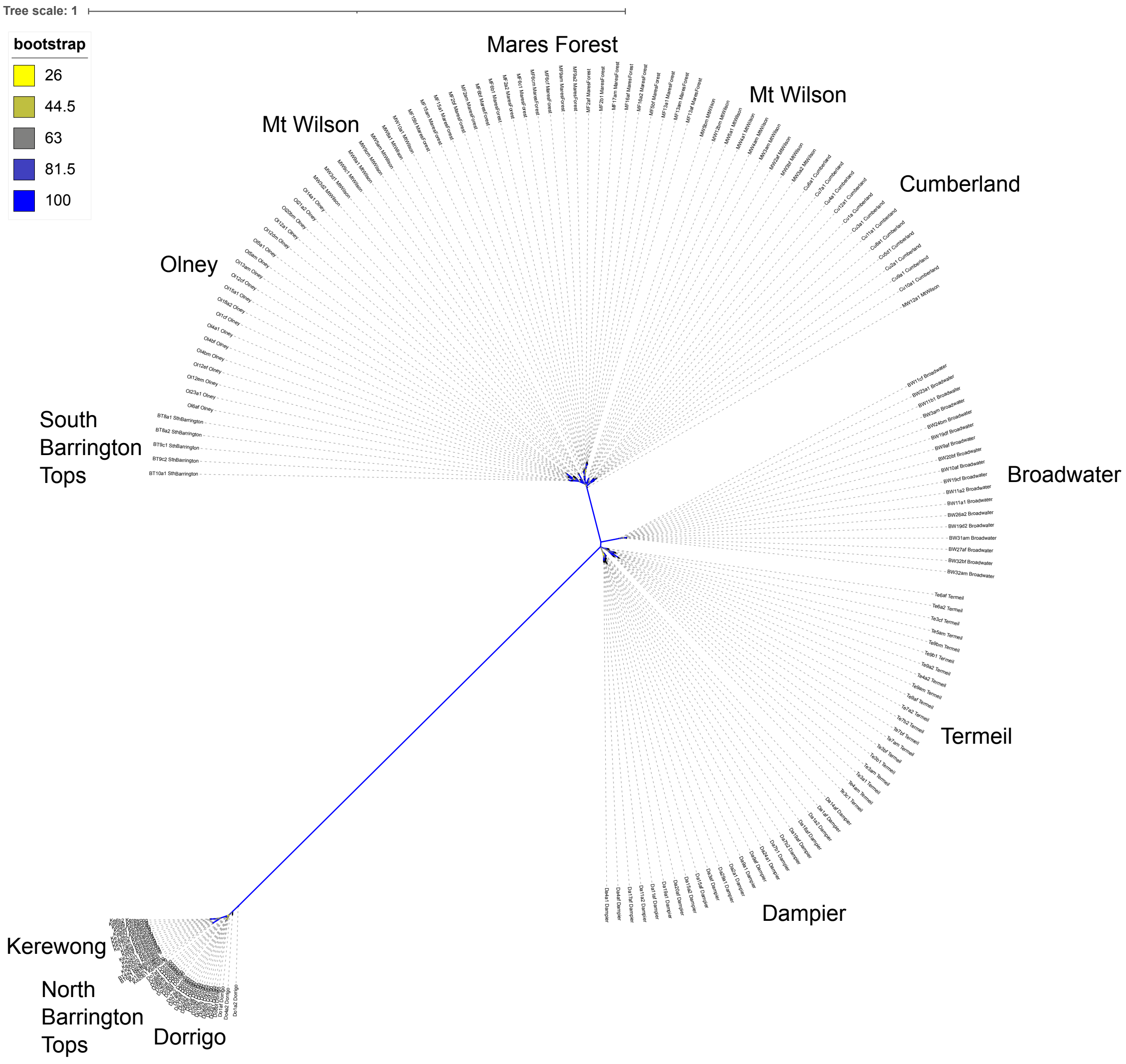


**Figure S2**. Maximum likelihood unrooted phylogeny inferred from 452 SNPs across for all individuals populations of *Austroplatypus incompertus*. Bootstrap support is coloured on the branches but indicates node support.

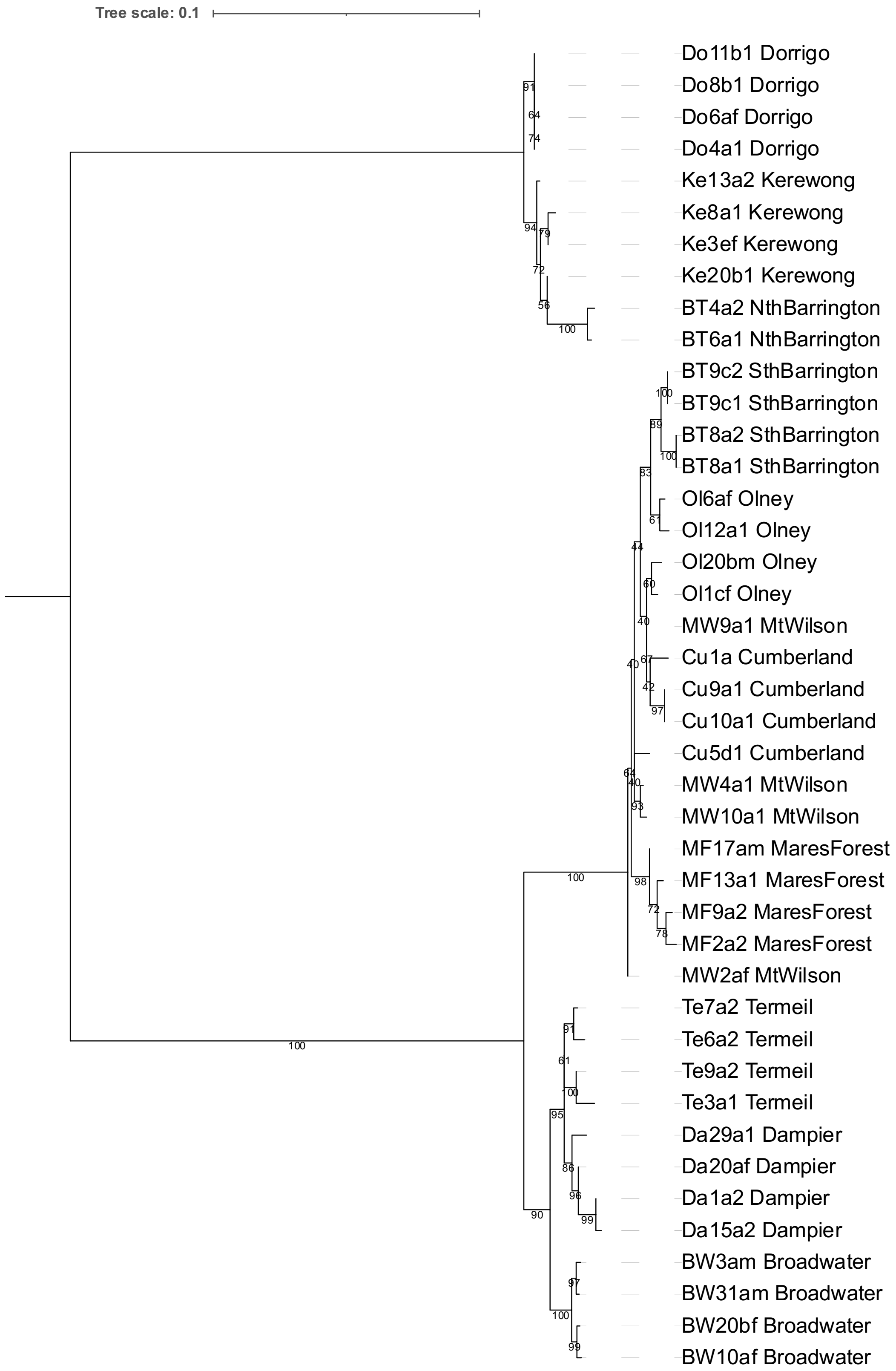


**Figure S3**. Maximum likelihood unrooted phylogeny inferred from 452 SNPs for four individuals per populations of *Austroplatypus incompertus*. Bootstrap support is given on the branches.

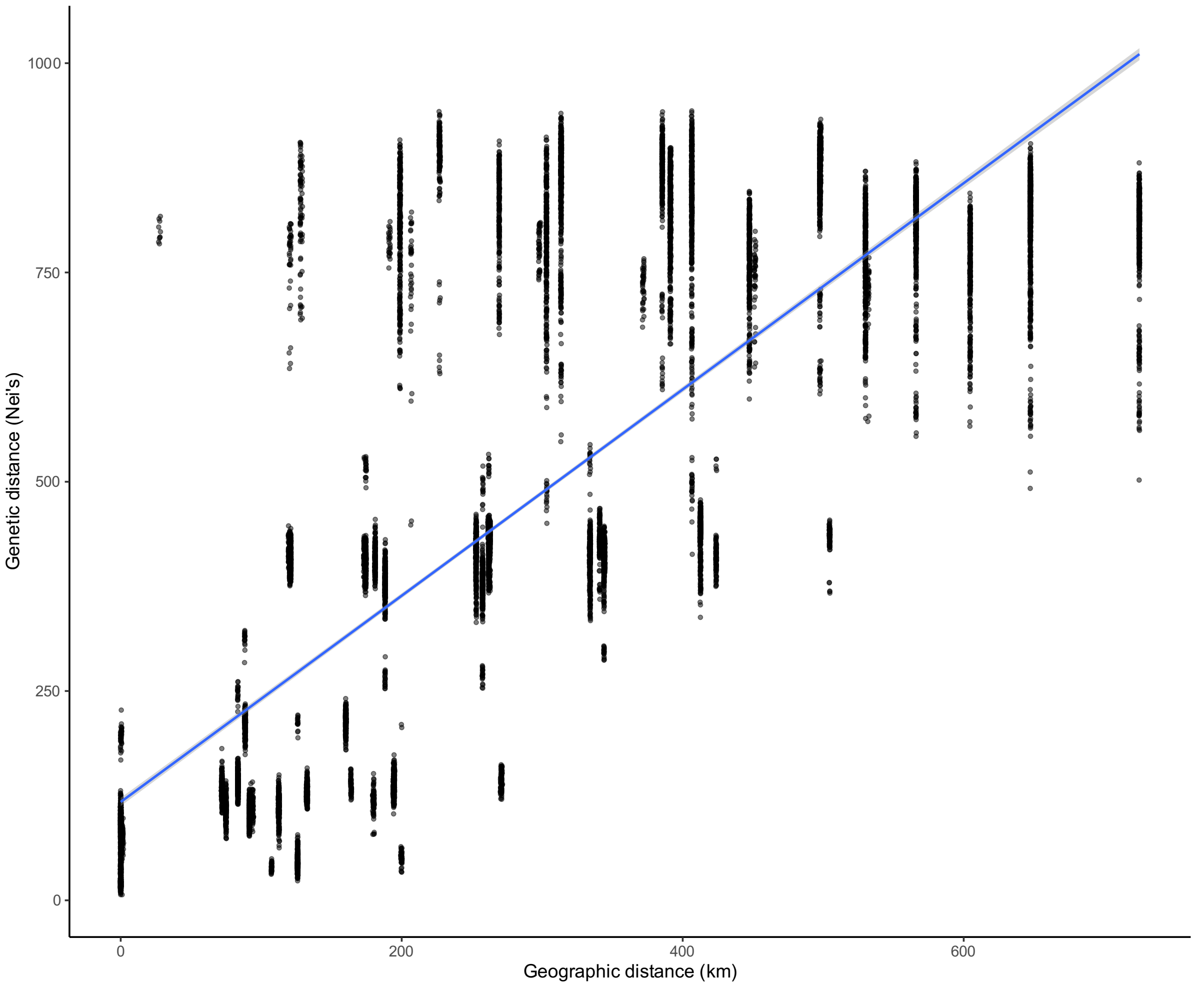


**Figure S4.** Linear modelling of pairwise genetic and geographic distances shows strong isolation by distance (IBD), with three predominant clusters of pairs of individuals separated by genetic and geographic distances. Geographic distances between individuals explain 62% of the variation of the observed genetic distances.
